# The impact of Relative Language Distance on Bilingual Language Control – a functional imaging study

**DOI:** 10.1101/771212

**Authors:** Keerthi Ramanujan

## Abstract

Cross-linguistic activation is unavoidable in bilinguals and they require language control to manage it. In this study, it is posited that Relative Language Distance (RLD; the extent of lexical feature-similarity between bilinguals’ languages) can affect the extent of cross-linguistic activation and hence influence bilingual language control. This was investigated via an er-fMRI word-translation task on three similar bilingual groups but with varying RLDs: Dutch-English (low-RLD), Hindi-English (intermediate-RLD) and Cantonese-English (high-RLD). Cross-linguistic conflict and the degree of conflict monitoring/control necessary to manage it were expected to *increase* with *decreasing* RLD across groups and be appropriately reflected in the activity of conflict monitoring/control neural regions, such as the ACC (anterior cingulate cortex). Analysis revealed a significantly differential ACC response across the groups, reflecting its adaptation to differential conflict monitoring/control demands generated by RLD. The findings provide emerging evidence for RLD as a dimension of bilingualism impacting bilingual language control processes and neurobiology.

## Introduction

How bilinguals juggle their two languages has been an area of intensive study in bilingualism research. Their evident success as skilled dual-language communicators is attributable to Language Control – cognitive processes that facilitate unhindered use of a desired target language while managing any potential language interference that might arise (Abutalebi & Green, 2007; Green, 1998). Such dual-language control is afforded via the Bilingual Language Control (BLC) network, comprising the prefrontal, inferior parietal and anterior cingulate cortices, the basal ganglia and the cerebellum (Abutalebi & Green, 2007, 2016; Calabria et al., 2018; Hervais-Adelman et al., 2011; Higby et al., 2013).

### Bilingual Heterogeneity & Adaptive Bilingual Language Control

While bilinguals, who make up half the world’s population (Grosjean & Li, 2013), have a common need for dual-language control, their dual-language experiences are certainly different and distinctive. Bilinguals hail from different sociolinguistic environments and vary along key dimensions such as Age of Bilingualism Onset, extents of their L1-L2 Immersion, Usage and Proficiency (Luk & Bialystok, 2013), as well as in Interactional Language Modes (single-language use vs. code-mixing, see Green, 2011, Green & Abutalebi, 2013). Such differences in bilingual experience are of significance because they have been shown to give rise to meaningful variation in bilingual neurobiology, particularly with respect to the BLC network (e.g., Berken et al., 2016; DeLuca et al., 2019; Perani et al., 1998, 2003; Pliatsikas et al., 2017; Tu et al., 2015). Acknowledging this variation, the Adaptive Control (AC) Hypothesis (Green & Abutalebi, 2013) proposes that heterogenous experiences of bilingualism could generate varying language control needs. To meet such varying needs, language control processes as well as the neural system implementing them undergo adaptation. Thus, adaptive effects/changes can manifest within the BLC network and may be observable as changes in either regional responsivity (functional activity changes), network connectivity/efficiency, or structural capacity (grey/white matter changes) (Green and Abutalebi, 2013). By positing the notion of neuroplastic adaptation, the AC hypothesis can therefore explain and predict neurobiological variation among bilinguals (Bialystok, 2017; Green & Abutalebi, 2013).

### Relative Language Distance & Bilingual Heterogeneity

Of the several dimensions that contribute to heterogeneity in bilingual experience and neurobiology, an understudied one concerns their languages themselves – specifically, the language pairings/combinations they speak. Variations in L1-L2 combinations are after all natural and pervasive among bilinguals. For instance, some bilinguals speak two relatively ‘close’ languages (e.g., Catalan and Spanish; Cantonese and Mandarin), while others are fluent in two ‘distant’ languages (e.g., Cantonese and English). Thus, across bilinguals, there is an obvious variation in the *relative* extents to which their two languages share linguistic features, at the lexical level (whole words, the phonemic repertoire, orthography/script) and at the morphosyntactic level (word order, subject-verb agreements, verb inflections etc) (Cenoz, 2003; Tolentio & Tokowicz, 2011). In the present study, this property of relative typological language similarity/difference of a spoken language pair is referred to as *Relative Language Distance* (RLD). Thus, bilinguals can differ in the dimension of L1-L2 RLD. Languages that are linguistically ‘closer’ often share evolutionary histories and as a result have noticeable similarities in perceptible lexical-level features such as script and phonology (e.g., Dutch and English, both Indo-European Germanic languages) compared to those that are linguistically ‘distant’ (e.g., Sino-Tibetan Mandarin vs. Dutch) (Cenoz, 2003). The question that arises now is, does this dimension of bilingualism matter for bilingual neurobiology? More specifically, would RLD have any impact on BLC processes and neurobiology the way other dimensions (such as AoA, proficiency etc) do? This is the focus of the present study.

### The Present Study: Relative Language Distance & Adaptive Bilingual Language Control

The underlying reason for bilinguals needing language control is the phenomenon of joint language activation. Inevitable co-activation of L1 and L2 representations during communicative processes such as language production renders them prone to cross-linguistic conflict and interference. These need to be efficiently controlled and managed for meaningful communicative output. While dual-language activation has been known to occur regardless of typological/RLD differences between language-pairs (see Degani et al., 2018 for a brief overview), it is possible that the *extent* of cross-linguistic joint activation could vary with the extent of relative similarity between L1 and L2’s perceptible linguistic features, i.e. the Relative Language Distance (RLD). When RLD of a given L1-L2 pair is *low* (e.g., Catalan-Spanish), cross-linguistic activation may be more extensive, with multiple lexical candidates across L1 and L2 (e.g., homographs, homophones, orthographic and phonological neighbours) being co-activated owing to the greater degree of ambiguity among similar L1 and L2 representations. However, when RLD is *high* (e.g., Tamil-English, Cantonese-English), extent of cross-linguistic activation may be much more reduced, potentially even limited to either L1 or L2, given the greater distinctness and language-specificity of L1 and L2’s lexical representations (Degani et al., 2018; Gollan et al., 1997; Dijkstra and Van Heuven, 2002). The normally language-*independent* cross-linguistic activation might thus be transformed into a language-*restricted* one as bilinguals’ RLD increases (for other instances of such ‘language-selectivity’, see Kroll et al., 2006).

Such an RLD-mediated effect can have implications for language control. Along with variation in the extent of cross-linguistic activation due to RLD, one would also expect variation in the extent of the ensuing cross-linguistic *conflict*. If bilinguals with varying RLDs experience varying magnitudes of cross-linguistic conflict, then, as per the AC Hypothesis, they would need different extents of language control to manage said conflict. Thus, lower the RLD (i.e. greater the similarity) of a language pair, greater the need for language control, especially cross-linguistic conflict/interference monitoring and control (see also Abutalebi et al., 2015 and Costa et al., 2006 for a similar argument). Therefore, the BLC network should undergo adaptation to meet the different conflict monitoring and control demands generated by variation in RLD.

To explore this possible effect of RLD on the BLC network, a functional neuroimaging study of word translation was carried out on three bilingual groups, each representing distinct levels of RLD (operationalized here as the extent of script and phonological similarity between L1 and L2 lexical representations^1^). The groups were:

i. High Distance Cantonese – English (hd-CE) bilinguals: This group’s two languages have visually distinct scripts (traditional Chinese characters vs Latin alphabets) with no overlapping orthographic representations. Considering that Cantonese is a Sino-Tibetan tonal language and English is a non-tonal Indo-European one, there is also little phonological similarity between the two (Zhou et al., 2010). This places the CE bilinguals at the higher end of the RLD continuum.
ii. Low Distance Dutch– English (ld-DE) bilinguals: Not only do lexical representations of both West-Germanic languages have high orthographic similarity (use identical Latin letters), they also have a relatively higher phonological overlap (Schepens et al., 2013)., making DE bilinguals a low distance group.
iii. Intermediate Distance Hindi – English (id-HE) bilinguals: Both languages have distinct orthographic representations (Devanagari vs. Latin script), with some phonological overlap (Sunderman & Priya, 2012). Compared to the Cantonese-English pair there is a greater phonological overlap due to shared Indo-European membership but certainly lesser than Dutch-English (Hindi: Indo-Iranian; English: Germanic). This places the HE group between the CE and DE groups with respect to RLD.

A printed word translation task was deployed over alternatives like picture-naming or lexical decision. Since the bilinguals in this study were also biliterate, a visual word translation task would tap into all three levels of lexical representation (orthographical, phonological and semantic). Moreover, translation requires that two languages – the ‘source’ (language of the translation stimulus) and the ‘target’ (output language) – be activated in rapid succession on *every* trial (de Groot & Christoffel’s, 2006; Kroll et al., 2018). This surely calls for the deployment of language control processes, specifically conflict monitoring processes, so that any potential interference from competing co-activated representations may be detected and resolved appropriately. The Anterior Cingulate Cortex (ACC)’s role as the key neural region implementing conflict monitoring/detection during linguistic and non-linguistic processes alike has been well established (Abutalebi et al., 2012; Botvinick et al., 1999, 2004; Branzi et al., 2016; MacDonald et al., 2000; van Heuvan et al., 2008). Since the groups would need to manage different levels of cross-linguistic conflict during word translation due to varying RLDs, adaptive effects manifesting as *regional responsivity differences* in the ACC were expected. Specifically, a comparatively greater ACC activation for low distance DE and lower ACC activation for high distance CE group was expected, while id-HE’s ACC activity was expected to be intermediate to DE and CE’s during word translation.

## Materials and Methods

### Participants

All participants were right-handed, neurologically normal biliterate bilinguals. They were included in the study on the basis of being highly proficient in their L1 (Cantonese/Hindi/Dutch) as well as L2 (English) and having adequate exposure and experience using both languages in their everyday lives. The three groups had very similar bilingual profiles and differed only with respect to the RLD of their L1-L2 pairs.

The CE group (n=19; 10 females; mean age = 21.95±1.78) comprised native Hong Kong-ers. The HE (n=19; 10 females; mean age = 21.37±2.27) and DE (n=20; 9 females; mean age = 21.75±2.07) groups comprised non-locals who at the time of the study were studying or working in Hong Kong. All three groups had comparable age, years of education and family income levels (see Table 1). Following Surrain & Luk (2017)’s recommendations, additional details on the groups’ sociolinguistic backgrounds and bilingual histories are provided in the Supplementary Materials to provide a clearer picture of their bilingual experiences. The study was approved by the Human Research Ethics Committee of the University of Hong Kong. Informed consent was obtained from all participants.

**Table 1:** Demographic Characteristics

|  | N(F/M) | AGE | Years of Education | Income Level |
| --- | --- | --- | --- | --- |
| *hd-CE* | 19 (10F/9M) | 21.95 (1.78) | 16.16 (0.9) | 6 |
| *id-HE* | 19 (10F/9M) | 21.37 (2.27) | 15.16 (1.80) | 5 |
| *ld-DE* | 20 (9F/11M) | 21.75 (2.07) | 16.05 (1.19) | 6 |
| p-value |  | 0.678 | 0.05 | 0.579 |

Table lists number of participants in each group along with mean Age, mean Number of Years of Formal Education (standard deviation in parentheses) and mode Income Level (participants selected an income range on a 6-point scale that best matched their family's total income during ages 5-18), along with p-values for one-way ANOVA on Age and Education Level and Kruskal-Wallis H Test on Income level.

### Language Proficiency Assessment

Participants provided subjective proficiency estimates for their L1 and L2 (English). Objective proficiency in L1 and L2 was assessed in two ways for each participant: (i). L1 and L2 naming accuracy on a 30-item timed picture naming task; (ii). Forward (L1→L2) and Backward (L2→L1) written translation accuracy for a total of 90 L1 and 90 L2 words in high, medium and low frequency categories. Pictures and translation stimuli for assessing L2 proficiency was identical across groups. The translation items for DE were free of interlingual homographs and cognates. Table 2 lists each group’s mean L2-English-AoA along with self-rated subjective and objective proficiency scores.

**Table 2:** L1 and English proficiency

|  | L2-AoA | Proficiency (self-rated) |  | Picture Naming<br>(max = 30) |  | Translation (max = 90) |  |
| --- | --- | --- | --- | --- | --- | --- | --- |
|  |  | L1 | L2 | L1 | L2 | L1→L2 | L2→L1 |
| *hd-CE* | 4.16 (1.98) | 9.01 (0.97) | 8.49 (0.68) | 28.05 (1.35) | 27.16 (1.34) | 84.58 (3.25) | 87.47 (3.86) |
| *id-HE* | 4.16 (1.54) | 8.59 (0.75) | 8.72 (0.53) | 26.16 (2.69) | 27.84 (1.30) | 84.21 (2.97) | 84.37 (3.09) |
| *ld-DE* | 6.70 (1.87) | 8.95 (0.86) | 8.31 (0.56) | 27.20 (2.93) | 26.95 (1.64) | 86.85 (3.52) | 87.15 (3.76) |

### fMRI experiment

The study comprised a printed-word translation task (TRAN) involving translation of L1 words (TRAN-L1 trials) and L2-English words (TRAN-E trials). A simple task of word reading (READ) involving reading L1 words (READ-L1 trials) and L2-English words (READ-E) which wouldn’t engage language control to the extent translation would (Price et al., 1999), served as an active baseline control.

#### Stimuli

TRAN and READ tasks each comprised 60 L1 (Cantonese/Hindi/Dutch) and 60 L2 (English) high familiarity, high frequency word stimuli with unambiguous translations (4 languages × 2 tasks = 8 word-lists). A one-way ANOVA followed by post-hoc Tukey-HSD revealed no significant differences (*p*>*0*.*01)* in mean word frequency^2^ across all stimuli lists (see Supplementary Materials for TRAN and READ stimuli’s corpora sources). Words did not repeat across the lists. Verbs and adverbs in all 4 languages were absent. Dutch-English homographs were avoided.

#### Experimental Design

Every participant performed TRAN and READ tasks in the following order: TRAN (Run1)-READ (Run1)-TRAN (Run2)-READ (Run2). Run2 was a repeat of Run1. Each participant read/translated 120 stimuli (60L1+60L2-English) in one run.

Both READ and TRAN tasks were implemented as pseudo-randomized language switching paradigms comprising “switches” (input language would change) and “stays” (input language remains the same), similar to Abutalebi et al., 2007 & 2012. This would also prevent any habituation effects that might otherwise arise from highly predictable switching patterns and, in this case, also from sustained unidirectional translation (see Chee et al., 2003; Grill-Spector et al., 2006).

There were equal number of switches and stays in each run (= 59). Their order along with order of L1 and L2-English words was pre-randomized for both runs and counterbalanced across all participants. There was a maximum of 4 stay trials before a switch trial for both tasks.

#### Procedure

For READ task, participants simply read out the displayed words. For TRAN task, they were instructed to translate L1 words to L2 (English), and identical L2 words into their respective L1 (Cantonese/Hindi/Dutch). Participants were previously familiarised with the translation task and were instructed to try and produce a translation for all words seen. This was to ensure that they would atleast activate the appropriate lexical representations even if they couldn’t articulate it quickly enough. They were also trained to verbalize their responses during the tasks in a manner that minimized head, jaw and tongue movements to limit motion-related imaging artefacts. The stimuli were displayed on a monitor (640×480 pixels, 4:3 aspect ratio) and presented through a mirror mounted on the head-coil. All stimuli (including fixation) were displayed as white fonts (size 36) on a black background (see Figure 1). TRAN run stimuli were displayed for 950ms while READ run stimuli were displayed for 750ms after which a fixation point – “ * “ – appeared. The duration of this fixation was jittered for either 1617, 3011 or 4613ms (Dale 1999). Technical constraints prevented the recording of in-scanner responses. However, a de-briefing session immediately followed the scanner session where participants assessed their in-scanner performance.

**Figure 1:**
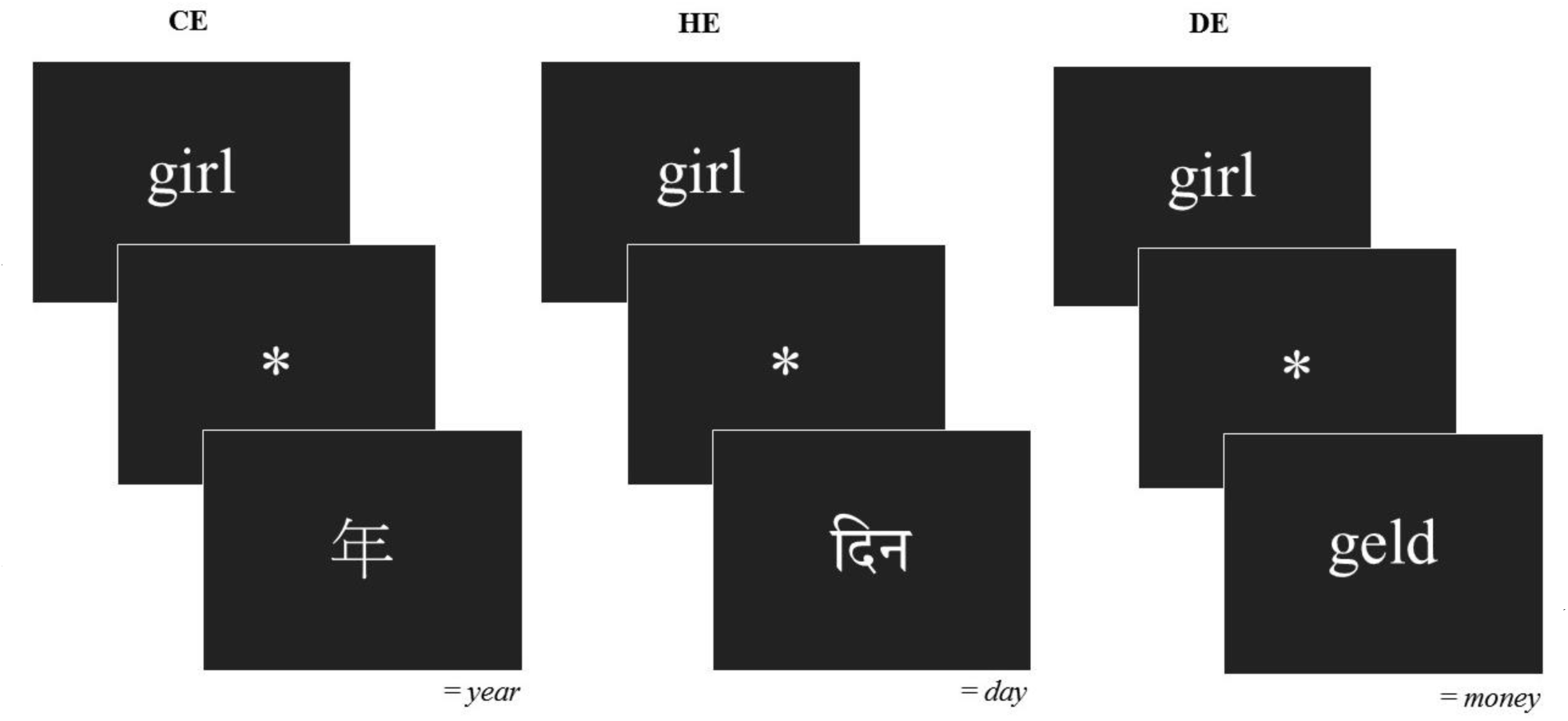
Examples of stimuli presented to CE, HE and DE bilinguals during TRAN/READ (words are depicted in larger font for clarity). The following ‘serif’ fonts were used to maintain visual uniformity between L1-English fontfaces: PMingLiU (Traditional Cantonese Characters); Sanskrit Text (Devanagari Hindi alphabet); Times New Roman (Latin alphabets).

#### MR Imge acquisition

All participants were scanned in a 3T Philips Achieva scanner (Philips Medical Systems, Best, NL) at the MR Imaging Unit of the University of Hong Kong.

Functional volumes for TRAN and READ were acquired using the same Echo Planar Imaging (EPI) sequence (Repetition Time [TR] = 2000 ms, Echo Time [TE] = 30 ms, Flip Angle = 90°, Field of View [FOV] = 240×240, matrix = 160 × 160, slice thickness = 2.5 mm, number of slices = 38 axial, voxel size = 1.5 × 1.5 × 2.5). To optimize the EPI signal, 10 dummy volumes preceded each run.

High resolution T1-weighted images were acquired using an MPRAGE (Magnetization Prepared Rapid Gradient Echo) sequence (150 slices, TR=8.03 ms, TE=4.1 ms; flip angle = 8°, voxel size = 1mm3 isotropic).

### fMRI data preprocessing and analysis

Functional volumes were preprocessed and analysed using SPM12 (v6906) (Wellcome Department of Cognitive Neurology, London, UK) running on Matlab2013b.

#### Preprocessing

The following standard preprocessing steps were performed for TRAN and READ tasks: 1). Slice-time correction using sinc-interpolation with the middle (19^th^) slice as reference; 2). Realignment using within-subject 6-parameter rigid body spatial transformation; 3). Unified Segmentation of T1 structural images. For CE, the ICBM space template for East Asian brains was used in the affine regularization step. For the HE and DE groups, the ICBM European brain template was used instead; 4). Normalization into standard MNI space by applying forward deformation fields derived from segmentation and resampling into 1.5 × 1.5 × 2.5 mm voxels; 5). Smoothing with a 6mm FWHM (full width at half-maximum) Gaussian kernel. The TRAN run of one DE participant and the READ run of another was discarded due to excessive head motion (>3mm in x, y, z direction). This resulted in 19 TRAN and READ functional data sets for the DE group, equal to the CE and HE group.

#### fMRI Statistical Analysis

Within group and between group effects of READ and TRAN tasks were assessed using the standard General Linear Model based two-level random effects (RFX) approach. At the first/subject level, two design matrices, one each for READ and TRAN tasks, were specified. For READ, trials of reading L1 (READ-L1) and L2-English words (READ-E) from each run were modelled in the design matrix. For TRAN, trials corresponding to translating L1 words (TRAN-L1) and L2-English words (TRAN-E) from each run were entered into the design matrix. Delta functions of these conditions of interest were convolved with the canonical hemodynamic response function (HRF) to create regressors for each condition. Additionally, the six movement parameters for each run were entered as covariates of no interest. A high pass filter with a 128s cut-off was used to remove scanner drifts. Serial correlations were treated using the AR (1) model.

At the second level, a 3 × 2 × 2 factorial model with factors of RLD (hd-CE id-HE, ld-DE), task (READ & TRAN) and input languages (L1 and L2-English) was created and estimated.

The following contrasts were assessed at the 2^nd^ level:

i. An F-contrast of TRAN to test for any RLD-induced activity differences across the groups’ BLC regions during the entire translation task, regardless of input language.
ii. An F-contrast of TRAN-E to test for any RLD-induced activity differences across the groups’ BLC regions during translation of identical L2-English stimuli.
iii. T-contrasts of TRAN>READ and TRAN-E>READ-E for the three groups to detect any increases in activation during translation relative to reading within their BLC networks.
iv. An F-contrast of READ to check for main effects of RLD within the BLC network, during L1 and L2 reading.

### ROI analysis and BOLD-Behavioural Correlational Analysis

Since we predicted ACC to show differential activity across the groups during translation, a Region of Interest (ROI) analysis was carried out using MarsBaR (Brett et al., 2002). The ROI was generated based on the peak ACC MNI co-ordinates reported in Abutalebi et al., 2012 where a conflict effect had been detected for bilinguals and monolinguals alike. Thus, an 8mm sphere centred around x=4, y=16, z=44 was created. Beta values/effect sizes were extracted from the ROI for each single-subject TRAN-E and TRAN-L1 contrasts. A one-way ANOVA followed by post-hoc testing using Tukey’s HSD was performed on combined beta values of TRAN-E and TRAN-L1 (overall TRAN) and on TRAN-E alone to test for differences in ACC-ROI activity across the groups.

Since the TRAN-E trails comprised identical stimuli for all three groups, a simple correlation and regression analysis was carried out between the groups’ BOLD activity in the groups’ ACC-ROI during TRAN-E and their L2-English language measures, viz., L2-English AoA, Self-rated L2-English Proficiency and Objective L2-English Proficiency (i.e., L2-English picture naming and L2-English word translation test scores).

## Results

### Language Assessment

One-way ANOVA with post-hoc testing on mean L2-English-AoA revealed that DE’s mean L2-English-AoA was significantly different from both HE and CE’s (*F*_*2,55*_ *=* 12.99, *p* < .01, *η*^*2*^_*p*_ = 0.32). However, comparisons of English proficiency revealed no significant differences (*p* > .01) between the groups for: i). self-rated English proficiency (*F*_*2,55*_ *=* 2.34, *p =* .1, *η*^*2*^_*p*_ =.08) or performance on ii). English picture naming (*F*_*2,55*_ *=* 2.02, *p =* .14, *η*^*2*^_*p*_ =.07) and iii). backward translation tasks (*F*_*2,55*_ *=* 4.31, *p =* .02, *η*^*2*^_*p*_ = .14). There were no significant differences amongst the groups in forward translation (*F*_*2,55*_ *=* 3.76, *p =* .03, *η*^*2*^_*p*_ = .12) as well. (See Figure 2). Pairwise T-tests revealed no significant L1-L2 differences (*p* > .01) for each group in: i). self-reported L1 and L2 proficiency and ii). L1 and L2picture naming performance, (See Figure 2). The forward and backward translation scores were not significantly different for HE and DE but differed significantly for the CE group (*t* = −3.55, *p* = .002). However, when translation accuracy of only High and Medium frequency words was considered, the differences went away. This was true for HE and DE as well.

**Figure 2:**
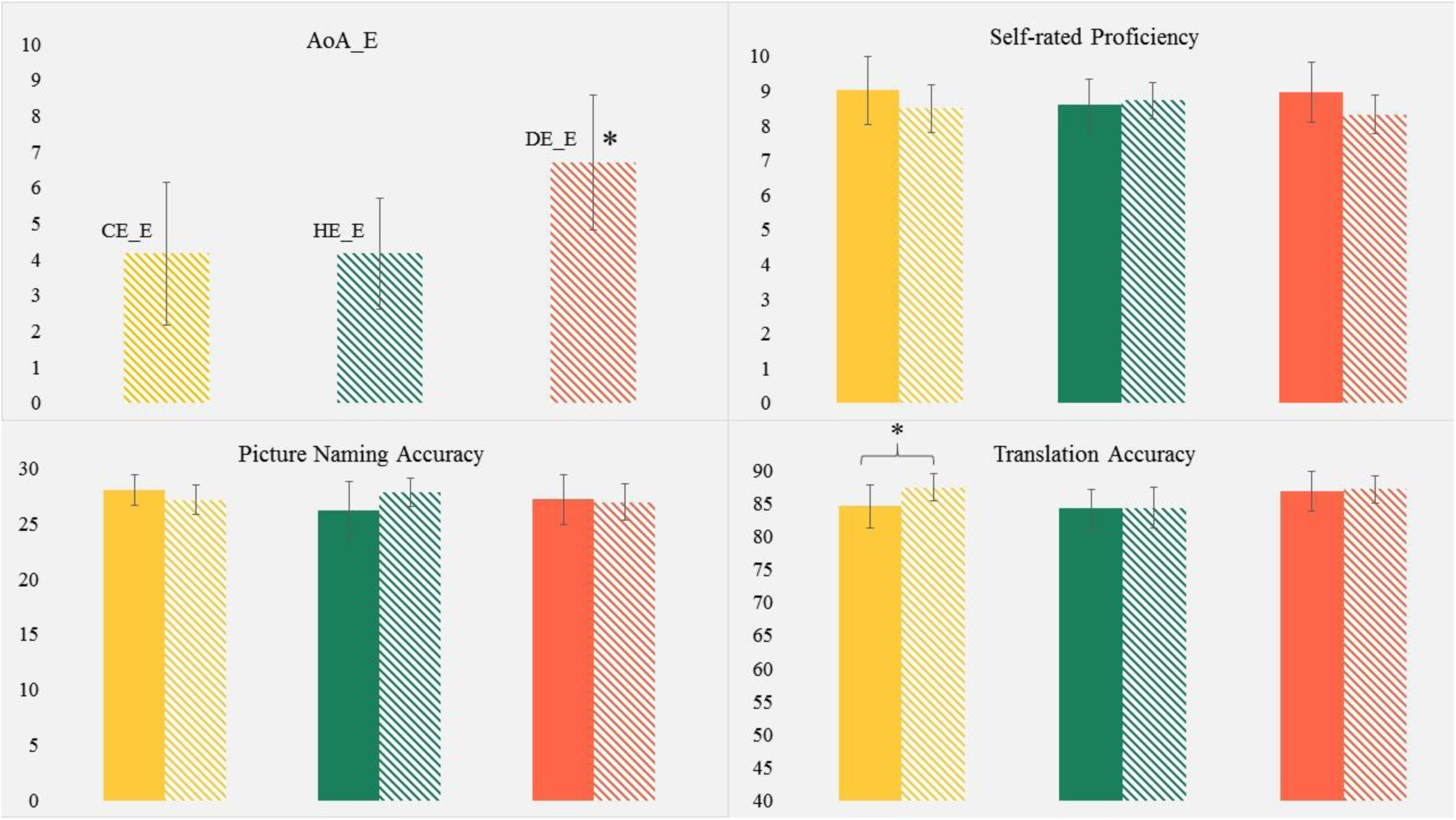
Comparison of CE (*yellow*), HE (*green*) and DE (*orange*) bilinguals’ Age of acquisition of English (AoA_E), Self-rated proficiency in L1 (*solid colours*) and L2-English (*stripes*), accuracy in translating L1 words (*solid colours*) and L2-English words (*stripes*) and performance during picture naming in L1 (*solid colours*) and L2-English (*stripes*). Error bars represent SD. No significant within-group and between-group differences were found, unless indicated with a ‘*’. See **Table 2** for exact values.

### Functional imaging

All participants reported high familiarity with all READ and TRAN stimuli in the de-briefing session. For READ, participants uniformly reported ceiling level accuracy. For TRAN runs, participants reported skipping no more than 8 words across both runs in each language resulting in a >=90% hit rate per language for each participant.

The results of the tested contrasts were considered significant if they survived a (uncorrected) voxel level threshold of *p* < .005 or lower along with cluster-level FWE (family-wise error) correction for multiple comparisons of *p* < .05. A spatial threshold of *k*≥100 contiguous voxels was applied.

i. The F-contrast testing for differential activity among the groups during TRAN task revealed significant differences in the Left Superior Temporal Gyrus (STG), Bilateral Rolandic Operculum and the ACC (see Table 3). The results at the ACC (x =12, y = 16, z = 40) survived an even more stringent threshold of FWE p < .05 at the *voxel* level (peak pFWE-corr = .008, k = 15 voxels).
ii. The F-contrast testing for RLD-generated differences across the groups during translation of identical L2-English words (TRAN-E), revealed significant differential activity in a large ACC cluster and in the right Rolandic operculum and the left Insula between the groups (see Table 3). The contrast estimates plotted at the ACC clusters revealed increased BOLD response for ld-DE compared to id-HE and hd-CE in both TRAN and TRAN-E conditions (See Figure 3).
iii. Only DE bilinguals showed significant ACC activation (x = 9, y = 22, z = 40) for TRAN>READ T-contrast. For the same contrast, HE showed a nearly significant activation of the cingulum at x = 8, y = −13, z = 38, (cluster *p*FWE-corr = .06). No suprathreshold clusters were discovered for CE at *p* = .005 or lower thresholds. For details on TRAN>READ T-contrast for each group, see Supplementary Materials 3a. For the TRAN-E>READ-E T-contrasts search volume limited to the right cingulate cortex (defined using an AAL template mask) revealed significant activation in the ACC (x =11, y = 31, z = 33) only for DE. Full list of TRAN-E>READ-E T-contrast activations for each group see Supplementary Materials 3b.
iv. Finally, F-Contrast for READ showed no significant differential activation due to RLD in the BLC regions across the groups.

**Table 3:** Main effect of Relative Language Distance across Groups

| Region | MNI co-ordinates (mm) |  |  | z-score | Peak p<br>(uncorr) | Cluster p<br>(FWE-corr) | k |
| --- | --- | --- | --- | --- | --- | --- | --- |
|  | x | y | z |  |  |  |  |
| **Translation (TRAN)** (cluster FWE $p<.05$) | | | | | | | |
| **R Rolandic operculum (BA 48)** | **59** | **-7** | **15** | **5.42** | **<.001** | **<.001** | **652** |
| R Rolandic operculum (BA 48) | 42 | 7 | 15 | 4.75 | <.001 |  |  |
| R Rolandic operculum (BA 48) | 50 | -9 | 15 | 4.25 | <.001 |  |  |
| **R Anterior/Mid Cingulate (BA 32)** | **12** | **16** | **40** | **5.23** | **<.001** | **<.001** | **325** |
| R Anterior Cingulate (BA 24) | 5 | 19 | 25 | 4.27 | <.001 |  |  |
| R Anterior/Mid Cingulate (BA 32) | 6 | 23 | 35 | 4.13 | <.001 |  |  |
| **R Superior Frontal (BA 6)** | **21** | **-1** | **45** | **4.46** | **<.001** | **<.001** | **271** |
| R Mid Cingulate (BA 32) | 9 | -4 | 43 | 4.28 | <.001 |  |  |
| **L Rolandic operculum (BA 48)** | **-44** | **-12** | **23** | **5.01** | **<.001** | **.001** | **255** |
| L Precentral (BA 48) | -51 | -4 | 23 | 3.87 | <.001 |  |  |
| L Superior temporal (BA 48) | -59 | -12 | 13 | 3.71 | <.001 |  |  |
| **Translation of only L2-English words (TRAN-E)** (cluster FWE $p<.05$) | | | | | | | |
| **R Rolandic operculum (BA 48)** | **59** | **-7** | **15** | **4.26** | **<.005** | **<.001** | **644** |
| R Rolandic operculum (BA 48) | 48 | -7 | 13 | 3.92 | <.005 |  |  |
| R Insula (BA 48) | 41 | 5 | 13 | 3.84 | <.005 |  |  |
| **R Anterior/Mid Cingulate (BA 32)** | **12** | **17** | **38** | **4.16** | **<.005** | **.013** | **307** |
| R Anterior/Mid Cingulate (BA 32) | 6 | 23 | 35 | 3.73 | <.005 |  |  |
| R Anterior Cingulate (BA 24) | 5 | 19 | 25 | 3.46 | <.005 |  |  |
| **L Insula (BA 48)** | **-41** | **-12** | **23** | **4.10** | **<.005** | **.009** | **326** |
| L Insula (BA 48) | -30 | -15 | 20 | 3.71 | <.005 |  |  |

Table shows clusters with minimum of 200 voxels per cluster > 8.0 mm apart; Bold Rows indicate peak voxels within cluster; Labels derived from AAL toolbox (Automatic Anatomical Labelling, Tzourio-Mazoyer et al., 2002) and Brodmann template provided in MRICron (Rorden & Brett, 2000)

**Figure 3:**
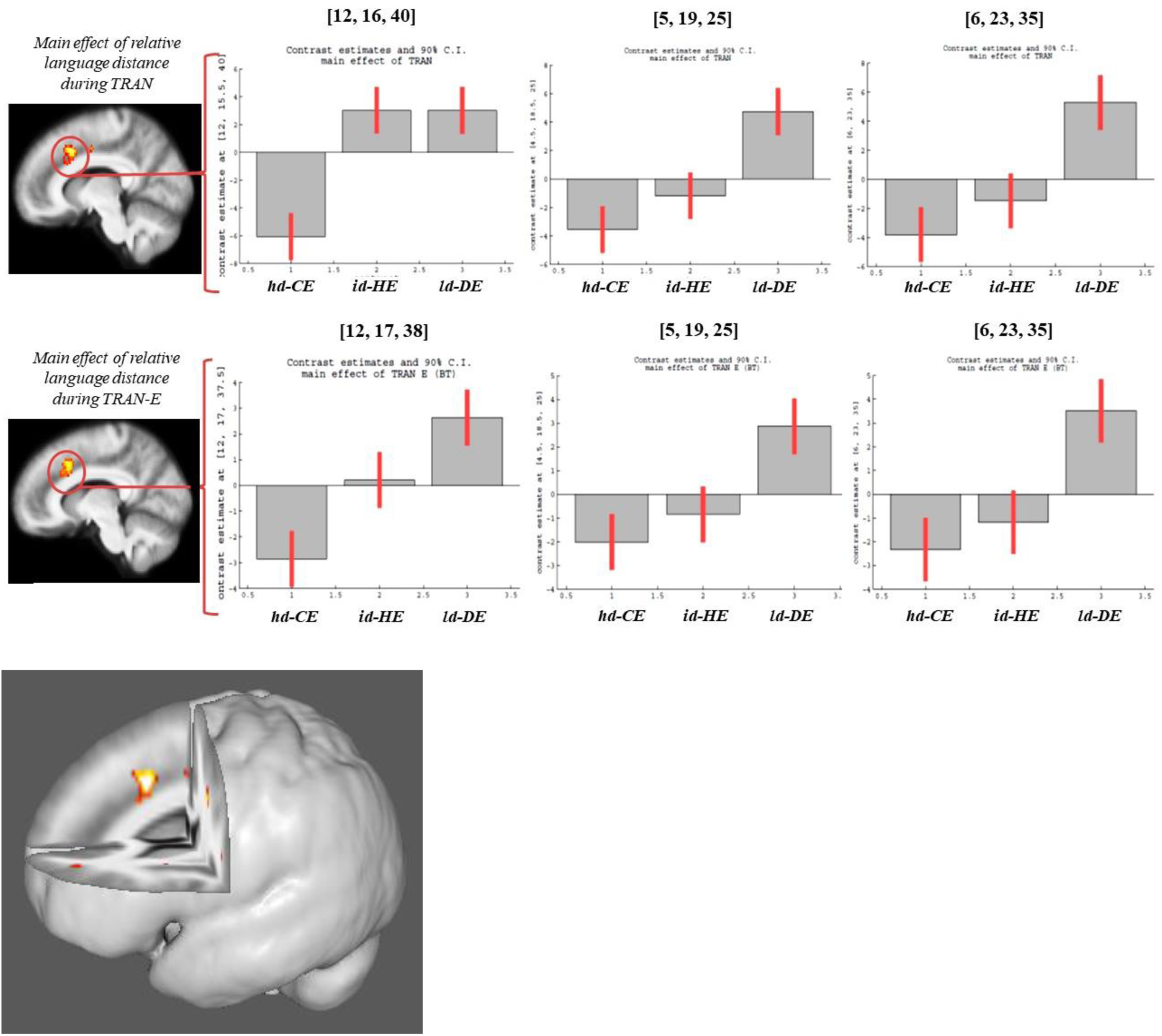
Main effect of RLD observable as significant differential activity (FWE corrected) in the ACC during the entire TRAN task (*top panel*) and specifically the TRAN-E trials (*bottom panel*) overlaid on mean structural image of the three bilingual groups, along with the contrast estimates plotted at voxels of the same ACC cluster. Bottom-most image depicts the same differentially activated ACC cluster during the TRAN task overlaid on a 3D version of the same mean structural image for clarity.

### ROI Analysis and BOLD-Behavioural Correlational Analysis

The one-way ANOVA on the extracted beta values from the ACC ROI indicated significant differences between the groups for TRAN (forward and backward translation combined) (*F*_*2,111*_ = 6.46, *p* = .002, *η*^*2*^_*p*_ = .10) and for TRAN-E (backward translation only) (*F*_*2,53*_ = 4.59, *p* = .01, *η*^*2*^_*p*_ = .15) (See Figure 4). Post-hoc testing showed that BOLD signal at the ACC ROI for the DE group was significantly higher than CE’s for TRAN and for TRAN-E (see Table 4).

**Table 4:** Post-hoc comparisons of BOLD response at ACC [4, 16, 44] 8mm spherical ROI

|  |  | TRAN (TRAN-L1 + TRAN-E) |  | TRAN-E only |  |
| --- | --- | --- | --- | --- | --- |
| | | Mean Difference | $p_{\text{Tukey}}$ | Mean Difference | $p_{\text{Tukey}}$ |
| CE | HE | -0.73 | 0.03 | -0.84 | 0.13 |
|  | DE | -0.98 | **0.002** | -1.28 | **0.01** |
| HE | DE | -0.25 | 0.65 | -0.44 | 0.55 |

Groups in the first column have been compared to groups in the second column.

**Figure 4:**
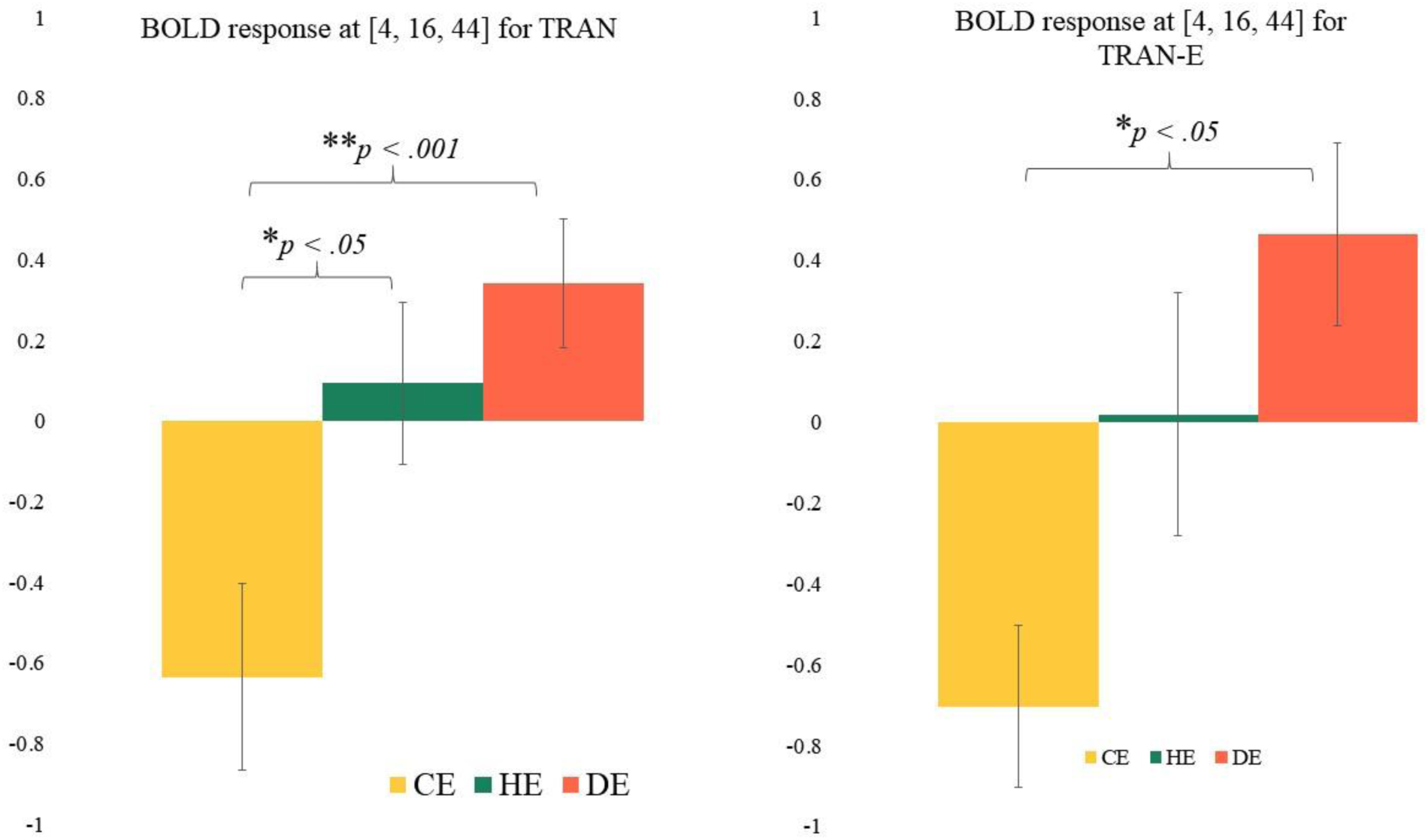
Effect sizes with for BOLD response at 8mm ACC ROI centred at *x* = 4, *y* = 16, *z* = 44 for (*left)* the entire Translation task (TRAN) and (*right*) for the L2-English translation trials (TRAN-E) only. Error bars represent SE.

ACC activity during TRAN-E did not significantly correlate (*p* > .05) with L2AoA, L2 proficiency or Backward Translation accuracy for any of the groups (see Figure 5) nor when the groups were collapsed (see Supplementary Materials, section 4).

**Figure 5:**
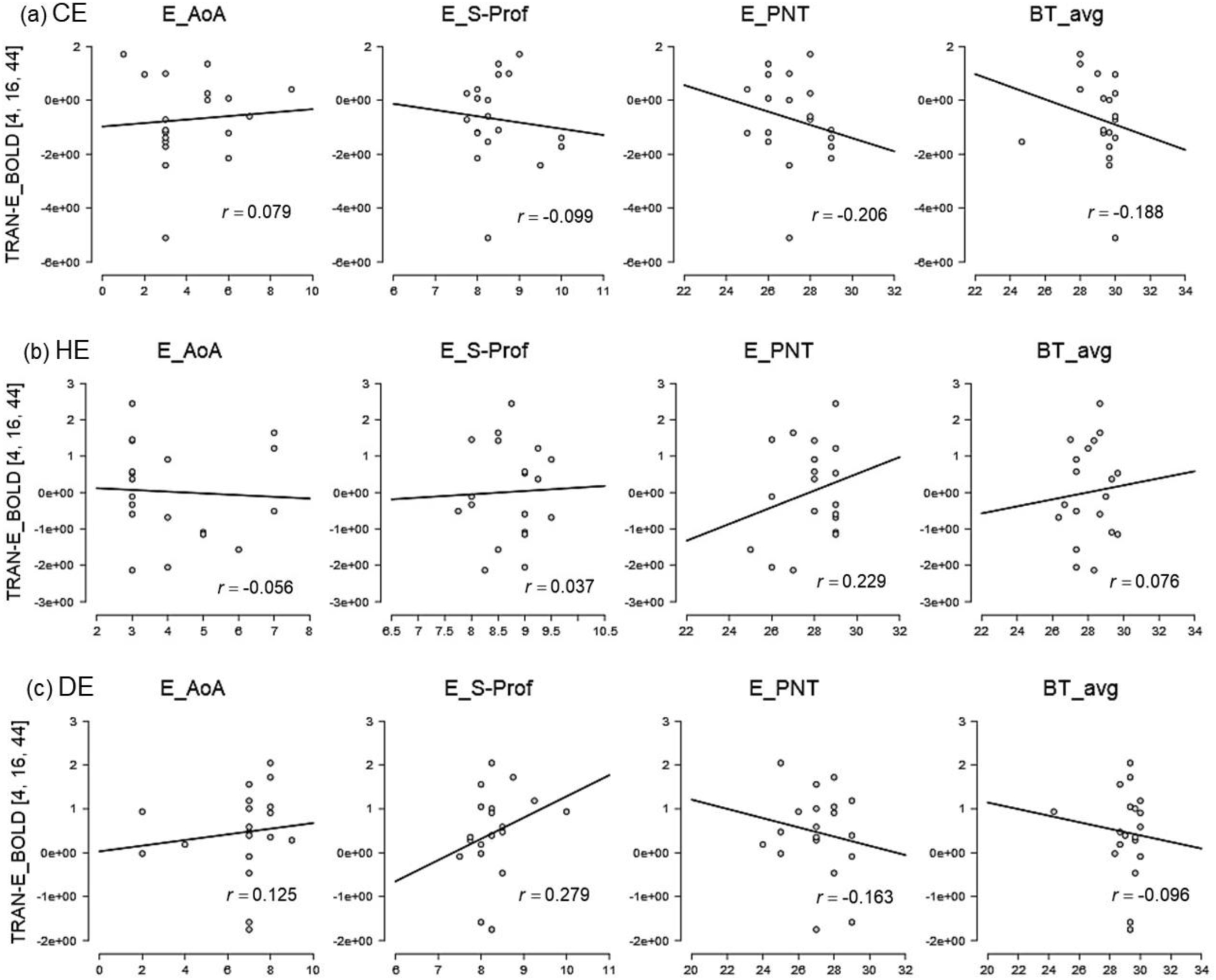
Group-wise correlations plots of ACC ROI’s activity during TRAN-E (y-axis) and L2 measures (x-axis), viz., L2-English AoA (“E-AoA”), Self-rated L2-English Proficiency (“E_S-Prof”), L2-English Picture Naming Test scores (“E_PNT”) and average Backward Translation (L2-English→L1) accuracy (“BT_avg”) with the respective Pearson’s Correlation Coefficients (*r*). No correlation was significant (all *p* > .05).

## Discussion

The Adaptive Control Hypothesis (Green and Abutalebi, 2013) advances the notion that the BLC network will adapt to different demands on language control that may arise due to different bilingual experiences and contexts. Such adaptation may be achieved via alteration in the BLC network’s regional responsivity, structural capacity, and/or its connectivity/efficiency. It was posited in this study that one dimension of bilingual experience, Relative Language Distance (RLD), might also modulate language control demands in bilinguals and consequently induce adaptation within their BLC networks. Specifically, it was expected that conflict monitoring demands would vary as a function of RLD – closer the L1 and L2, greater the extent of cross-linguistic conflict generated and therefore, greater the degree of conflict monitoring and control required. As per the AC hypothesis, the differential conflict monitoring demands generated by varying RLDs should cause adaptation in the neural substrate responsible for conflict control, i.e. the ACC. To test this, an er-fMRI version of word translation (a linguistic task requiring conflict monitoring and control) was used to specifically elicit activity in the ACC and capture any task-dependant responsivity differences in the region. It was predicted that when bilinguals with varying RLDs perform said word translation, functional responsivity in the groups’ ACCs would vary inversely with increasing RLD, i.e. in the pattern of ld-DE > id-HE > hd-CE.

The neuroimaging results seem to confirm this expectation. There was a main effect of RLD on conflict monitoring activity across the groups. This was observed as a significant differential activation of the groups’ ACCs during translation (Figure 3 & Table 3). ld-DE showed a significantly greater involvement of the ACC compared to id-HE and hd-CE groups during the task. The results indicate that variation in RLD resulted in differential engagement of the conflict management region (ACC) for an identical linguistic task.

The TRAN-E (L2-English → L1) results further attest the observed RLD-mediated effect on ACC activity. If RLD were to be inconsequential for adaptive bilingual language control and the network implementing it, then one might expect little-to-no significant difference in ACC neural activity when three similar, high proficient bilingual groups translated identical L2-English words into their respective L1s. Yet this was not the case. The groups’ ACC activity varied even during the task of translating identical semantic, phonological and orthographic representations (common English stimuli). Moreover, the ACC activity of each group during TRAN-E was not confounded by their L2 proficiency levels, L2 AoA or Backward (L2→L1) Translation ability as evidenced by the lack of correlation between ACC response and these measures (Figure 5 & Supplementary Materials 4). This is not surprising. Relative Language proficiency, which was anyway comparable across the groups of this study, might not have been able to alter the extent of language-independent co-activation or the magnitude of the ensuing cross-linguistic conflict in the present study, the way RLD, the critical factor the study’s bilinguals varied on, could (see also Abutalebi et al., 2013b for a similar conclusion regarding proficiency and ACC conflict control response). Such comparable relative language proficiency of the bilinguals in the study may also be the reason for the absence of significant differential activity in other BLC network regions such as the caudate, putamen, prefrontal and parietal cortices in this study. Indeed, the prefrontal cortex, the caudate and the putamen are known to be sensitive to relative language proficiency differences, exhibiting greater activity when bilinguals engage their less proficient language (e.g. Abutalebi et al, 2013a & b; Meschayan & Hernandez, 2006; Parker-Jones et al., 2012; Perani et al., 2003; Wang et al., 2007). Thus, in the present study, the lack of differences in these regions during translation, are not surprising. Similarly, there is no reason to expect differences in the parietal regions sub serving attentional control for the bilinguals performing translation. Considering these results together, the differential adaptive response exhibited by the ACC in this study can be attributed to the factor of relative language distance, rather than relative language proficiency or other factors. This further substantiates the notion of RLD driving adaptive functional responsivity changes in the ACC, via its influence on conflict monitoring needs. These results also re-affirm the ACC’s role as a reliable detector of cross-linguistic conflict, acting at the level of *local* language control, i.e. control of just the relevant subset of lexical representations, rather than entire set of L1/L2 lexical representations (i.e., *global* control) (see Branzi et al., 2016). Word translation can only be performed by necessarily accessing *both* languages (i.e., both source and target languages need to be ‘activated’), but only a limited set of lexico-semantic representations in either (the source stimulus, its semantic equivalent in the output language and any other co-activated potential competitors), needs monitoring and control.

There is substantial evidence of linguistic stimuli triggering language-independent/non-selective lexical co-activation in bilinguals that extends to all levels of representation – semantic, orthographic and phonological. L1 and L2’s phonological and semantic representations are known to coactivate in bilinguals with varying RLD levels (e.g., English & French: Jared & Kroll, 2001; Dutch & English: Lagrou et al., 2013; English & Welsh: Martin et al., 2009; Russian & English: Marian & Spivey, 2003; Japanese & English: Hoshino & Kroll, 2007; Hindi & English: Sunderman and Priya, 2012; Mandarin & English: Wu and Thierry, 2010). Shared orthographic representations between L1 and L2 further increases the spread of such joint activation (e.g., Bijeljac-Babic et al., 1997; Libben and Titone, 2009; Duyck et al., 2007; van Heuvan et al., 1998). However, when L1 and L2 orthographic representations are dissimilar in cases of high RLD, the obvious distinctness of their visual linguistic features (script) could limit the spread of such co-activation, restricting it to only those representational levels where there is some possible overlap, such as the phonological and/or semantic levels. Note that in the present study, to “level the playing field” as far as task stimuli were concerned, Dutch-English homographs were not used, since HE and CE would naturally not encounter them anyway due to higher RLD of their language pairs. Among the groups in the study, DE, by virtue of their low RLD, faced the greatest extent of cross-linguistic activation arising from the co-activation of phonological and orthographic candidates spanning *both* languages. For HE, the extent of co-activation would certainly not involve any orthographical representations due to the distance and disparity between Hindi and English. It could, however, be limited to any overlapping (and hence competing) phonological representations instead. Finally, for the CE group whose languages are the most distant, cross-language activation might be restricted to only the relevant L1 or L2 subsets as neither orthographic nor phonological lexical representations of their languages bear any resemblance to one another and therefore do not co-activate and co-compete during the translation process. Since each group varied in their requirement of conflict monitoring demands, the ACC as expected underwent adaptation, responding accordingly. Plots of contrast estimates from the ACC cluster revealed the exact directionality of its differential activation – higher activity in the ld-DE group, lower activity for the hd-CE and intermediate activity for the id-HE group for the TRAN and TRAN-E conditions (Figure 3). Comparison of the BOLD activation effect size at an independent ACC ROI further confirmed the same pattern of activation: ld-DE > id-HE > hd-CE (Figure 4). Taken together, these results confirm the assumptions regarding the inverse relationship between RLD, cross-linguistic conflict control demands and ACC response.

### Language Distance does matter for Bilingual Neurobiology and Language Control

The idea that cross-linguistic similarity/differences (however they might be referred to or operationalized) can impact linguistic neurocognitive processes and neurobiology is certainly not new (e.g., Bolger et al., 2005; Li, 2013; Li et al., 2014). Prior neuroimaging studies indicate that RLD certainly has some consequences for bilinguals, especially with regards to how each language is neurally represented and processed vis-a-vis the other. Linguistic features specific to one language can require additional or specialized neural resources (such as processing tonal vs. non-tonal phonemes), which can give rise to functionally different neural processing profiles for each language in the bilingual brain (e.g., Bick et al., 2011; Mei et al., 2015; Perfetti et al., 2007; Green et al., 2007; Tolentino & Tokowicz, 2011). However, despite language combinations/RLD being an obvious point of variation among bilinguals and a distinguishing dimension of the bilingual experience, its possible contribution to the bilingual experience has not received a lot of attention (Bialystok, 2017). The notion that RLD can influence the extent of co-activation in bilinguals finds mention and support in psycholinguistic models like the BIA+, which clarifies that non-selective lexical access in an integrated bilingual lexicon can be constrained by the extent of cross-linguistic overlap in orthographic, phonological and semantic codes across bilinguals’ two languages (see Dijkstra & Van Heuven, 2002, pp. 182-183). That RLD is also capable of influencing bilingual language control needs and in turn impacting underlying BLC network neurobiology is supported by neurocognitive models like the AC Hypothesis (Green & Abutalebi, 2013). By positing that idea that the underlying neural circuitry for language control can adapt to meet changing language control demands, the AC model implicitly allows for experiential dimensions of bilingualism such as proficiency, exposure, interactional modes and RLD to drive adaptive effects in the BLC network via their influence on language control needs.

There is now increasing evidence indicating that the degree of typological similarity/difference between bilinguals’ L1-L2 has a discernible impact on lexical processing (e.g., Casaponsa & Dunabeitia, 2016; Orfanidou & Sumner, 2005; Mosca et al., 2018), morpho-syntax processing (e.g., Tolentino & Tokowicz, 2011 and references therein; Momenian et al., 2018; Putnam et al., 2018), second and third language learning (e.g. Ghazi-Saidi & Ansaldo, 2017; Cenoz et al., 2003) and even executive functioning (e.g., Coderre and van Heuven, 2014b; van Heuvan et al., 2011; Yang et al., 2017). The focus of the present study was to explore whether such relative typological similarity/difference, or RLD as instantiated in this study, would have any effect at all on bilingual language control processes and network. The findings affirm that it does. It was possible to adequately isolate RLD’s effect by ensuring that the experimental groups varied only with respect to this feature but otherwise had very similar bilingual experiences, and were comparable on key dimensions of bilingualism such as proficiency and exposure that are already known to influence BLC neurobiology.

Although, the study provides emerging evidence concerning the influence of a less explored dimension of bilingualism on BLC neurobiology, it is not without its limitations. The evidence reported in this study specifically pertains to RLD-induced adaptive *responsivity* changes occurring in a task-dependant context in one hub of the BLC network (the ACC). It is not yet clear what other types of neural adaptations (functional connectivity, structural capacity changes etc.) RLD might engender within the BLC network. This is certainly an objective for future research. Further research is indeed required to establish and qualify RLD’s influence on the bilingual experience and how it might also interact with other key dimensions of bilingualism such as proficiency or exposure.

Not only is the prevalence of bilingualism continuing to increase, but so is the ease of access to diverse bilingual populations. At this time, it is important that bilingual heterogeneity and variability be acknowledged and the effect of differentiating experiential factors, such as RLD, on bilingual neurobiology and cognition be closely studied. This would help contribute to an even deeper understanding of the phenomenon of bilingualism.

## Acknowledgements

The support of the following individuals towards this study is gratefully acknowledged: Pasquale Della Rosa for his advice regarding neuroimaging experiment implementation; Linda Zhang for help with Cantonese and Eileen Waegemaekers for help with Dutch stimuli selection; The technical staff at the University of Hong Kong’s 3T MRI Unit in MRI data collection assistance; Davide Fedeli for assistance during initial neuroimaging analysis; and Jubin Abutalebi for his comments on an early version of this manuscript.

The study was supported by internal funding from The University of Hong Kong (HKU)’s and HKU Faculty of Education’s Faculty Research Fund (FRF) with additional support from HKU University Research Committee (URC) and the HKU Foundation for Educational Development and Research.

## Footnotes

1 As previously mentioned, RLD can also be defined with respect to morphosyntactic similarity between language pairs. But RLD was intentionally operationalized at the lexical level for the following reasons: i). bilingual language control mechanisms during lexical retrieval have been better studied compared to the more complex syntax-related processes; ii). the difficulty and practical challenges of controlling sentence-level variables across the four languages used in the present study.

2 Because linguistic differences do not permit written Cantonese and Hindi words to be matched with English words on features like orthographic neighbourhood density, word length and syllable count in the manner possible for Dutch and English words, all stimuli were matched on frequency.

## Notes

**Declaration of interest** The author declares that they have no known competing financial interests or personal relationships that could have appeared to influence the work reported in this paper.

