## Supplementary Materials for "The impact of Relative Language Distance on Bilingual Language Control – a functional imaging study"

1. Bilingual profile and history of study groups

Apart from knowing to speak, read/write their L1s (Cantonese/Hindi/Dutch) and L2-English, the participants, based on their responses in questionnaires and personal interviews, were also similar on the following socio-linguistic aspects:

i). CE, DE and HE members were all born and raised in Hong Kong, The Netherlands and northern India (e.g., Delhi) respectively and had been living in these countries in the past 2 years at the time of testing. All participants had therefore spent a significant part of their lives (childhood until end of formal schooling/pre-university) in regions where the L1 was the dominant and official indigenous language and English a very commonly used/learnt foreign L2 (English also enjoys official status in India and Hong Kong). The socio-economic and cultural climate of the countries/regions that the participants hailed from is such that being bilingual at a fairly early age, with English as the preferred L2 was quite common. Crucially, none of the participants had immigrant status in their respective home countries. This limited the number of languages they knew and used in familial and non-familial social circles to just the L1 and L2-English. In this manner, all participants had adequate opportunities for exposure and immersion in dual-language contexts.

ii). All participants across the groups groups reported formal study of both L1 and L2-English in school for atleast 6 years. Without exception, the medium of instruction and examination for all participants at the end of school and throughout undergraduate study (or higher, wherever applicable) was English.

iii). Most of the HE and DE participants were full-time students at the University of Hong Kong. Despite being away from their home countries they reported continued use of their L1 with friends and family via their real and virtual social networks. None were familiar with or learning Cantonese/Mandarin at the time of the study.

iv). All participants hailed from very similar socio-economic backgrounds (urban/semi-urban upbringing in single/double income households having 0-2 siblings and atleast one parent with college level education).

2. **Corpora sources for languages in study**

| | |
|---|---|
| *Dutch* | SUBTLEX-NL |
| *Hindi* | EMILLE-CIIL (monolingual written + spoken) |
| *Cantonese* | HK Corpus of Chinese Newspaper |
| *English* | SUBTLEX-EN(US) |

3a. **T-contrast TRAN>READ**

Voxel level threshold $p < .001$; Cluster-level FWE correction of $p < .05$; extent threshold, $k$=100 voxels

*Note:* For the CE group, no suprathreshold cluster were discovered at voxel level $p < .001$ or .005. Regions listed below for CE group alone are at threshold of $p < .01$

| Region | *MNI co-ordinates (mm)* | | | *z*-score | Cluster *p* (FWE-corr) | *k* |
|---|---|---|---|---|---|---|
| | *x* | *y* | *z* | | | |
| ***CE*** (@ *p* < .01) | | | | | | |
| **R Precuneus (BA 7)** | **17** | **-79** | **45** | **4.47** | **.025** | **580** |
| L Cuneus (BA 7/18) | -3 | -76 | 35 | 4.31 | | |
| L Superior Parietal (BA 19/7) | -18 | -81 | 45 | 3.37 | | |
| **R Supplementary Motor Area (BA 4)** | **12** | **-28** | **53** | **4.26** | **.001** | **1101** |
| L Precuneus (BA 4) | -6 | -36 | 58 | 3.89 | | |
| R Superior Parietal (BA 5) | 18 | -46 | 73 | 3.78 | | |
| ***HE*** (@ *p* < .001) | | | | | | |
| **R Superior Temporal (BA 22)** | **57** | **-49** | **20** | **5.34** | **<.001** | **479** |
| R Angular (BA 41) | 48 | -48 | 23 | 4.17 | | |
| R Mid Temporal (BA 21) | 45 | -52 | 15 | 3.86 | | |
| **R Postcentral (BA 3)** | **32** | **-37** | **53** | **5.27** | **<.001** | **727** |
| R Postcentral (BA 3) | 36 | -37 | 60 | 5.16 | | |
| R Postcentral (BA 2/3) | 48 | -27 | 43 | 5.01 | | |
| **R Anterior/Mid Cingulate (BA 23)** | **8** | **-13** | **38** | **5.33** | **.061** | **137** |
| R Anterior Cingulate (BA 23) | 5 | -4 | 40 | 3.63 | | |
| R Mid Cingulate (BA 23) | 11 | -6 | 33 | 3.39 | | |
| **L Paracentral Lobule (BA 4)** | **-5** | **-34** | **73** | **4.75** | **<.001** | **481** |
| R Supplementary Motor Area (BA 4) | 3 | -22 | 58 | 4.72 | | |
| R Precuneus (BA 5) | 5 | -45 | 58 | 4.57 | | |
| **L Mid Temporal (BA 37)** | **-41** | **-57** | **13** | **4.15** | **.042** | **151** |
| L Mid Temporal (BA 37) | -53 | -63 | 8 | 4.10 | | |
| L Mid Temporal (BA 41/21) | -42 | -48 | 13 | 3.49 | | |
| **R Rolandic Operculum (BA 48)** | **42** | **-27** | **20** | **4.14** | **.066** | **134** |
| R Supramarginal (BA 48) | 50 | -21 | 25 | 3.53 | | |
| R Supramarginal (BA 48) | 47 | -28 | 30 | 3.26 | | |

| | | | | | | |
|---|---|---|---|---|---|---|
| ***DE*** (@ *p* < .001) | | | | | | |
| **R Supplementary Motor Area (BA 6)** | **17** | **11** | **65** | **5.33** | **.02** | **180** |
| R Supplementary Motor Area (BA 6) | 12 | 4 | 65 | 4.47 | | |
| **R Anterior Cingulate (BA 32)** | **9** | **22** | **40** | **5.00** | **.002** | **275** |
| R Anterior Cingulate (BA 32) | 9 | 31 | 35 | 4.55 | | |
| **R Insula (BA 48)** | **44** | **14** | **-3** | **4.87** | **<.001** | **508** |
| R Rolandic Operculum (BA 48) | 48 | -7 | 13 | 4.63 | | |
| R Insula (BA 48) | 50 | 4 | -3 | 4.38 | | |
| **L Insula (BA 48)** | **-27** | **31** | **10** | **4.80** | **.03** | **164** |
| L Inferior Frontal (BA 47) | -32 | 28 | 0 | 3.98 | | |
| L Insula (BA 48) | -29 | 20 | 10 | 3.61 | | |

Table lists clusters greater than 100 voxels in size. Bold face indicates peak voxels within cluster.

3b. **T-contrast TRAN-E>READ-E (English word Translation > English word Reading)**

Contrast was thresholded as follows: Voxel level threshold *p* < .001; Cluster-level FWE correction of *p* <.05; extent threshold, *k*=10 voxels. Small volume correction (SVC) was then applied by limiting search volume using a predefined Right Anterior Cingulum mask (based on AAL template).
Voxel co-ordinates along with their peak *p* values, Z-scores and FWE-corrected cluster level *p* values are listed. SVC failed to find significant suprathreshold voxels for CE and HE groups in the right cingulate region but are nevertheless listed here for comparison with DE.

| Region | *MNI co-ordinates (mm)* | | | *z*-score | Cluster *p* (FWE-corr) | *k* |
|---|---|---|---|---|---|---|
| | *x* | *y* | *z* | | | |
| ***CE*** | | | | | | |
| R Mid Cingulum (BA 23) | 5 | -36 | 33 | 3.29 | .180 | 5 |
| ***HE*** | | | | | | |
| R Mid Cingulum (BA 23) | 11 | -13 | 35 | 3.63 | .146 | 8 |
| ***DE*** | | | | | | |
| **R Anterior/Mid Cingulum (BA 32)** | **11** | **31** | **33** | **3.73** | **.008** | **79** |
| R Anterior/Mid Cingulum (BA 32) | 11 | 23 | 35 | 3.61 | | |
| R Anterior/Mid Cingulum (BA 32) | 9 | 22 | 40 | 3.59 | | |
| R Anterior/Mid Cingulum (BA 32) | 12 | 26 | 33 | 3.53 | | |

4. Correlation of ACC ROI's activity during TRAN-E (translating L2-English → L1) with L2AoA, L2 proficiency measures and Backward Translation accuracy, collapsed across groups. No correlations were significant (all $p > .05$).

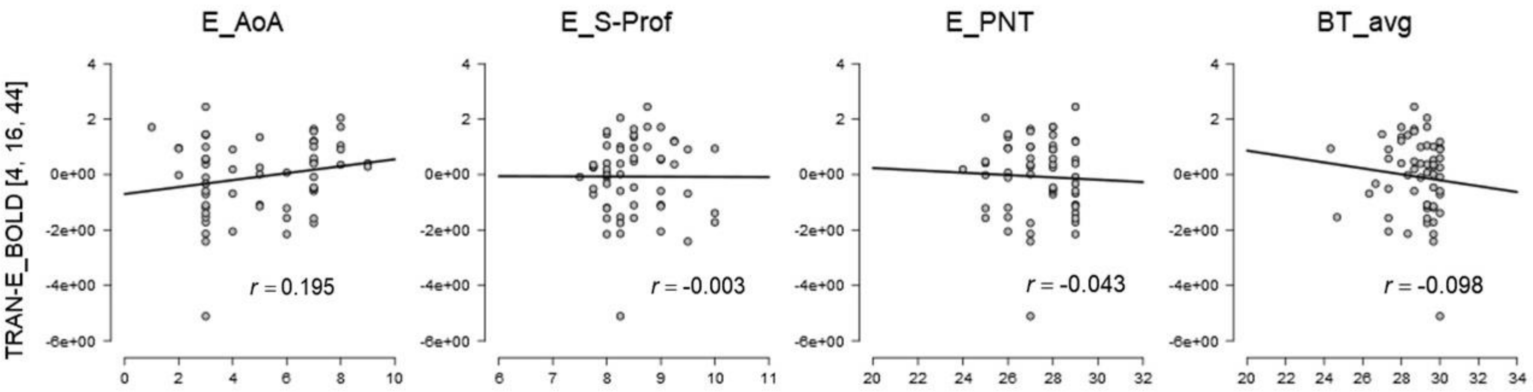


**Supplementary Figure1:** Correlation of ACC ROI's activity during TRAN-E (y-axis) of all 3 groups plotted against L2 measures (x-axis), viz., L2-English AoA ("E-AoA"), Self-rated L2-English Proficiency ("E_S-Prof"), L2-English Picture Naming Test scores ("E_PNT") and average Backward Translation (L2-English→L1) accuracy ("BT_avg") with the respective Pearson's Correlation Coefficients (*r*). No correlation was significant (all $p > .05$).
